# Title: Toleration of Frameshift Mutations in mRNA Sequences Encoding the N-terminal Peptides of Bacterial Type III Effectors

**DOI:** 10.1101/2023.04.10.536236

**Authors:** Jielin Yang, Moyang Lu, Mingyang Yu, Xinlong Wang, Ziyi Zhao, Lijun Luo, Xuxia Cai, Runhong Chen, Yueming Hu, Yejun Wang

## Abstract

Gram-negative bacteria deliver effector proteins into eukaryotic host cells through type III and type IV secretion systems, causing infections and diseases. It remains unclear about the signals guiding the specific secretion of the effectors. Here, we adopted an *in silico* approach to analyze the mRNA sequences encoding the putative peptides essential for effective secretion and translocation of type III and IV effectors. A surprisingly high proportion of type III effectors showed tolerance on frameshift mutations in signal-encoding mRNA sequences, and in contrast, very low percentage of type IV effectors showed the similar frameshift tolerance. The type III effectors with frameshift tolerance of secretion signals were widely distributed in effector or signal families and bacterial species. Natural frameshifts could be identified in type III effector genes, which were often remedied in time by nearby paired insertions or deletions. Frameshift-derived peptide sequences also retained the common properties present in the signal peptides of raw type III effectors. Natural language processing models were adopted to represent the common features in the mRNA sequences encoding N-terminal peptides of type III effectors or C-terminal peptides of type IV effectors, with which transfer learning models could well predict the effectors, especially type IV effectors. The observations in the study would facilitate us understand the nature and evolution of secretion signals of type III and IV effectors.

**Significance:** It has been a debate on the nature of signals for translocation of type III secreted effectors for a long time. Meanwhile, there has been no examination on the possibility of mRNA being as translocation signals for type IV or other types of secreted effectors. By computational simulation, the study demonstrated the protein nature of translocation signals for both type IV effectors and most type III effectors. Despite wide frameshift tolerance and atypical common features in mRNA sequences encoding the putative N-terminal signal sequences of type III effectors, more typical common physicochemical and amino acid composition properties between the mutation-derived and raw peptides, and the frequent self-correction phenomenon for naturally happening frameshifts supported the translocation signals at protein level of type III effectors. The common features in mRNA sequences encoding the translocation signal peptides of type III and IV effectors could also be combined in models for better prediction of the effectors respectively.

## Introduction

Gram-negative bacteria can manipulate type III or IV secretion systems to deliver substrate effectors into eukaryotic host cells, causing infections and pathogenicity (Hu et al., 2017; Galán and Waksman 2018). These effector proteins have large diversity among species, and are recognized by corresponding secretion systems through the signal sequences, which are located in the N-termini for type III effectors and the C-termini for type IV effectors (Lloyd et al., 2002; Nagai et al., 2005). The signal sequences of effectors secreted through distinct secretion systems contain specific atypical features, such as amino acid composition biases, physicochemical properties, evolutionary profiles, secondary structure, and so on (Arnold et al., 2009; Samudrala et al., 2009; Burstein et al., 2009). The signal features have been widely adopted in machine learning models to predict effectors accurately (Hui et al., 2020; Jing et al., 2021; Wagner et al., 2022; Wang et al., 2014; Noroy et al., 2019).

However, there have been debates on the nature of signal sequences of type III effectors (Anderson and Schneewind 1997; Mudgett et al., 2000; Rüssmann et al., 2002; Ramamurthi and Schneewind 2003; Niemann et al., 2013). Frameshifting mutations of mRNA sequences encoding the N-termini of *Yersinia yopQ* and many other genes did not abolish the secretion of the proteins through type III secretion conduits, but synonymous mutations on some codons blocked the secretion of some effectors, suggesting the mRNA nature of type III secretion signals for these effectors (Anderson and Schneewind 1997; Anderson et al., 1999a; Mudgett et al., 2000; Rüssmann et al., 2002; Ramamurthi and Schneewind 2003). A mechanism was proposed to explain the export of type III effectors, which coupled mRNA translation with the specific secretion of the encoded polypeptides (Anderson and Schneewind 1999b; Arnold et al., 2010). Another study found that the 25 nucleotides in 5’-untransltational regions (5’-UTR) of five genes encoding *Salmonella* effectors, *gtgA*, *cigR*, *gogB*, *sseL* and *steD*, were sufficient to mediate the translocation of Cya fusions into host cells in an Hfq-dependent manner (Niemann et al., 2013). In fact, other studies also disclosed the importance of noncoding or coding mRNA sequences in guiding protein secretion. For example, the 5’-UTR of *fliC* gene was sufficient to mediate the export of corresponding protein product by the flagellar system of *Escherichia coli* (Majander et al., 2005). The mRNA sequence encoding a C-terminal region of *Yersinia enterocolitica* YopR is required for the specific type III secretion of the effector protein (Blaylock et al., 2008). Translation-independent localization of mRNA in bacteria may provide explanations for the group of effectors whose translocation is mediated by the mRNA signals (Nevo-Dinur et al., 2011; Kannaiah and Amster-Choder 2014; Fei and Sharma 2018).

However, more evidence supports the signals are present in the N-terminal peptide sequences, since the significant changes of amino acid composition caused by mRNA frameshifts completely abolished the secretion of type III effectors (Lloyd et al., 2001; Karavolos et al., 2005). *In silico* modeling also suggested that only a few effectors could tolerate frameshifting mutations in mRNA encoding the N-terminal sequences (Arnold et al., 2009; Wang et al., 2011). However, there remained more than 10% effectors that could tolerate 5’-mRNA frameshifts (Wang et al., 2011). Since the computational models were based on the composition and/or physicochemical properties of amino acids in the N-terminal sequences, the effectors that tolerated 5’-frameshifts would have similar amino acid features with those of raw sequences. Consistently, proteins were frequently found with similar critical physicochemical properties before and after mRNA frameshifts (Bartonek et al., 2020; Wang et al., 2022). During the last more than ten years, many more new type III effectors have been validated, and the proportion of mutation tolerating effectors should be re-assessed to get a more general picture. Besides, there have been very few studies addressing the protein or mRNA nature for the signal sequences of type IV secreted effectors or other proteins, which should also be questioned. Finally, it would be interesting to observe whether there are common atypical signals buried in mRNA sequences, which can be used for recognition of the effector proteins.

## Results

### 1. Wide tolerance of frameshifts in mRNA encoding N-termini of type III effectors

We collected the 618 verified type III secreted effectors from TxSEdb, together with their encoding mRNA sequences (Methods). The N-terminal 100-aa peptide sequences were scanned by T3SEppML (Hui et al., 2020) with the default parameters, and 87.9% (543/618) were recalled correctly (Figure 1A,B; Supplemental Dataset S1). Frameshifting mutations were introduced to the 5’-ends of mRNA sequences causing frame changes (both “+1” and “-1”) of the whole effector proteins (Methods). The N-terminal 100-aa peptide sequences derived from the mutated nucleotide sequences were also predicted with T3SEppML. Surprisingly, among the effectors whose raw sequences were correctly recalled, 68.5% (372/543) showed tolerance of “+1” or “-1” frameshifts (Figure 1A,B; Supplemental Dataset S1). The percentage is much larger than previously estimated (Wang et al., 2011). To exclude the possibility that the observation of frameshift tolerance was caused by the false positive prediction by T3SEppML, we also choose another state-of-the-art model, Bastion3 (Wang et al., 2019), to repeat the analysis. Bastion3 showed a higher recalling rate, 90.1% (557/618); however, 82.9% (462/557) of the correctly recalled effectors could tolerate the frameshifts causing completely altered N-termini (Figure 1C; Supplemental Dataset S1).

**Figure 1.**
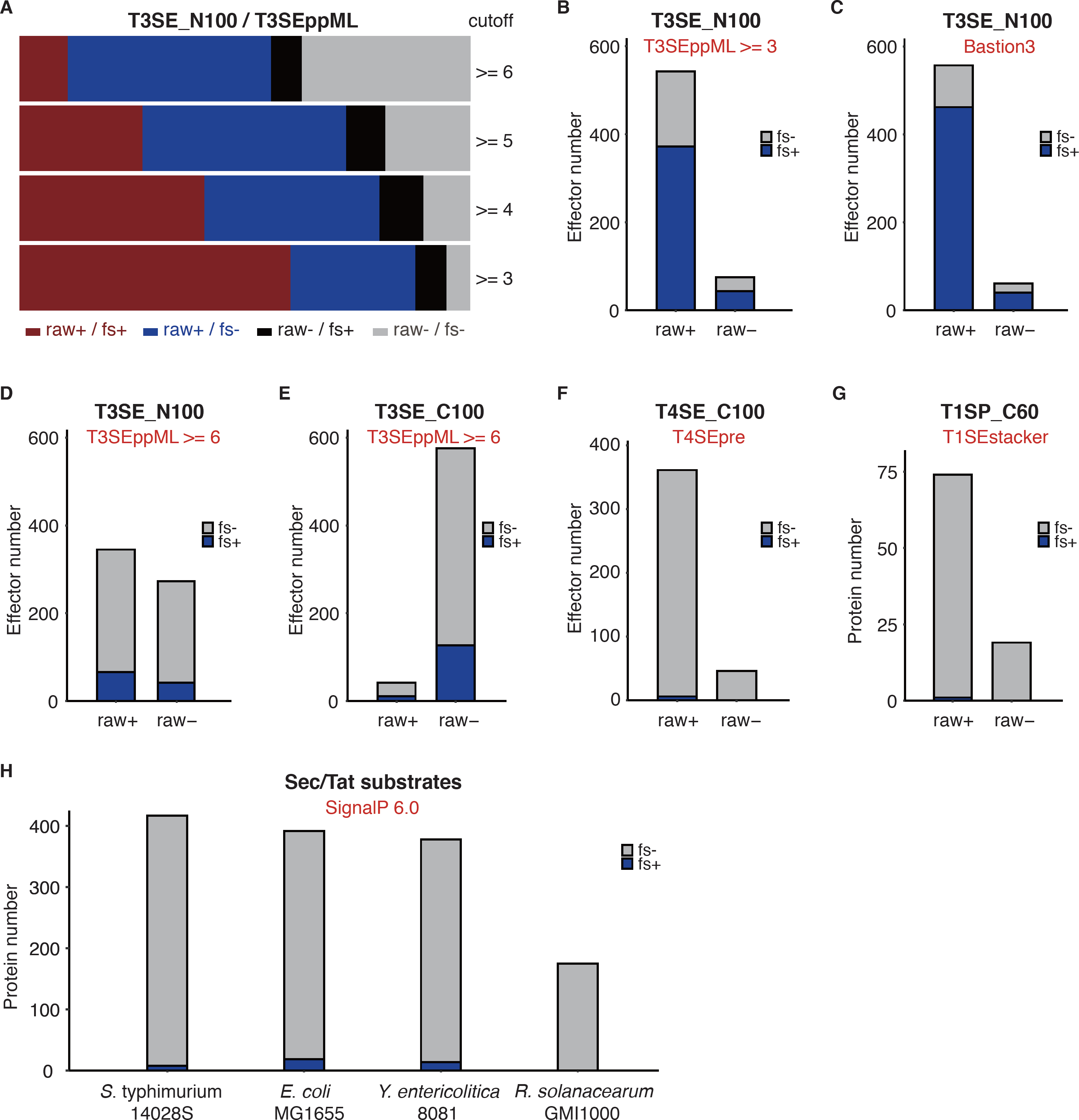
Tolerance of frameshifts in mRNA sequences encoding the signal peptides. **(A)** Distribution of the T3SEppML prediction results on the N-terminal 100-aa peptides of raw and frameshift-mutated type III effectors. Each aligned column in all the rows represented the same single effector. Each type III effector was classified as a combinatorial group of “raw+/fs+”, “raw+/fs-”, “raw-/fs+” or “raw-/fs-“, where “raw+” represented a raw sequence with positive prediction, “fs+” represented a frameshift-mutated sequence with positive prediction, “raw-” represented a raw sequence with negative prediction and “fs-” represented a frameshift-mutated sequence with negative prediction. **(B)** - **(H)** The distribution of secreted proteins or controls tolerant to the frameshift mutations in the nucleotide sequences encoding the putative signal peptides. The software tools and parameters used for prediction of the secreted proteins were also indicated. Default parameters were used if not specified.

We shifted the cutoff to the maximum for T3SEppML so as to reduce the false positive predictions at most. The recalling rate for the raw effectors was compromised to 55.8% (345/618), and the frameshift-tolerating rate also decreased apparently, to 19.1% (66/345) at the same setting (Figure 1A,D; Supplemental Dataset S1). As a control, the C-terminal 100-aa sequences for only 6.1% (31/509) of the raw effectors were predicted to contain type III secretion signals, and similar percentage of the raw positive predictions show tolerance of mRNA frameshifts (19.4%, 6/31) (Figure 1E; Supplemental Dataset S1). The recalling rates for N-termini and C-termini were significantly different (345/618 vs. 31/509; EBT, *p*=0.00), suggesting that the false positive rate was well controlled at this strict cutoff. In fact, the false positive rate could still be overestimated, since there were type III effectors with C-terminal signals (Tomalka et al., 2012; Login and Wolf-Watz 2015).

Taken together, the results demonstrated that a considerable proportion of type III effectors could tolerate mRNA frameshifts causing significant peptide sequence changes of the N-termini.

### 2. Low tolerance of type IV, type I and Sec/Tat secretion signals on mRNA frameshifts

To observe whether the frameshift tolerance is common for peptide-based signal sequences of various types of secreted proteins, similar observations were made for type IV effectors (C-terminal 100-aa sequences), type I secreted proteins (C-terminal 60-aa sequences), and Sec/Tat substrates (SignalP 6.0 positive predictions) (Methods). The raw and frameshifting peptide sequences were predicted with T4SEpre (Wang et al., 2014), T1SEstacker (Chen et al., 2022) and SignalP 6.0 (Teufel et al., 2022) respectively.

For type IV effectors, the recalling rate on the raw C-terminal sequences was 88.7% (361/407) at the largest specificity, and only 1.7% (6/361) of the positive results showed frameshift tolerance (Figure 1F; Supplemental Dataset S2). For type I secreted proteins, 79.6% (74/93) were correctly recalled based on the raw C-terminal peptide sequence, among which only 1.4% (1/74) tolerated mRNA frameshifts encoding the C-termini (Figure 1G; Supplemental Dataset S3). The Sec/Tat substrates were scanned from the whole genome of *Salmonella typhimurium* 14028S, *E. coli* MG1655, *Y. entericolitica* 8081 and *Ralstonia solanacearum* GMI1000 with SignalP 6.0, generating 392, 417, 378 and 0 candidates, respectively. Among these candidate Sec/Tat substrates, only 4.9% (19/392), 1.9% (8/417), 3.7% (14/378) and 0.0% showed tolerance on the mRNA frameshifts, respectively (Figure 1H; Supplemental Dataset S4).

Therefore, compared to type III effectors, the signal sequences of other types of secreted proteins in Gram-negative bacteria showed apparently low tolerance on mRNA frameshifts.

### 3. Family preference of type III effectors tolerating frameshifts in mRNA encoding the N-terminal sequences

With the 66 and 372 type III effectors whose N-terminal peptides contain secretion signals and tolerate mRNA frameshifts at the strict and relaxed cutoff respectively, we evaluated the distribution of effector or signal families (Hui et al., 2020), bacterial species and mutation patterns of tolerance (Supplemental Dataset S5). With the strict cutoff, only 17.5% (43/246) and 20.8% (55/265) of the recalled effectors showed frameshift tolerance, which were mapped to 35.9% (28/78) of the covered effector families and 26.2% (43/164) of the signal families, respectively (Figure 2A-B; Supplemental Dataset S5). None of the larger families (>=5 effectors covered) was significantly enriched with frameshift tolerant effectors (EBT, *p* > 0.05), although Effector_FAM_4 (4/10), Effector_FAM_11 (3/7), SigFAM_2 (3/7) and SigFAM_5 (3/6) showed relatively higher proportion of frameshift tolerance (Supplemental Dataset S5). We also observed the families with >= 2 covered effectors that all showed frameshift tolerance, and noticed three such effector and four signal families, Effector_FAM_46 (2/2), Effector_FAM_66 (2/2), Effector_FAM_73 (2/2), SigFAM_10 (3/3), SigFAM_32 (2/2), SigFAM_40 and SigFAM_43 (2/2) (Supplemental Dataset S5).

**Figure 2.**
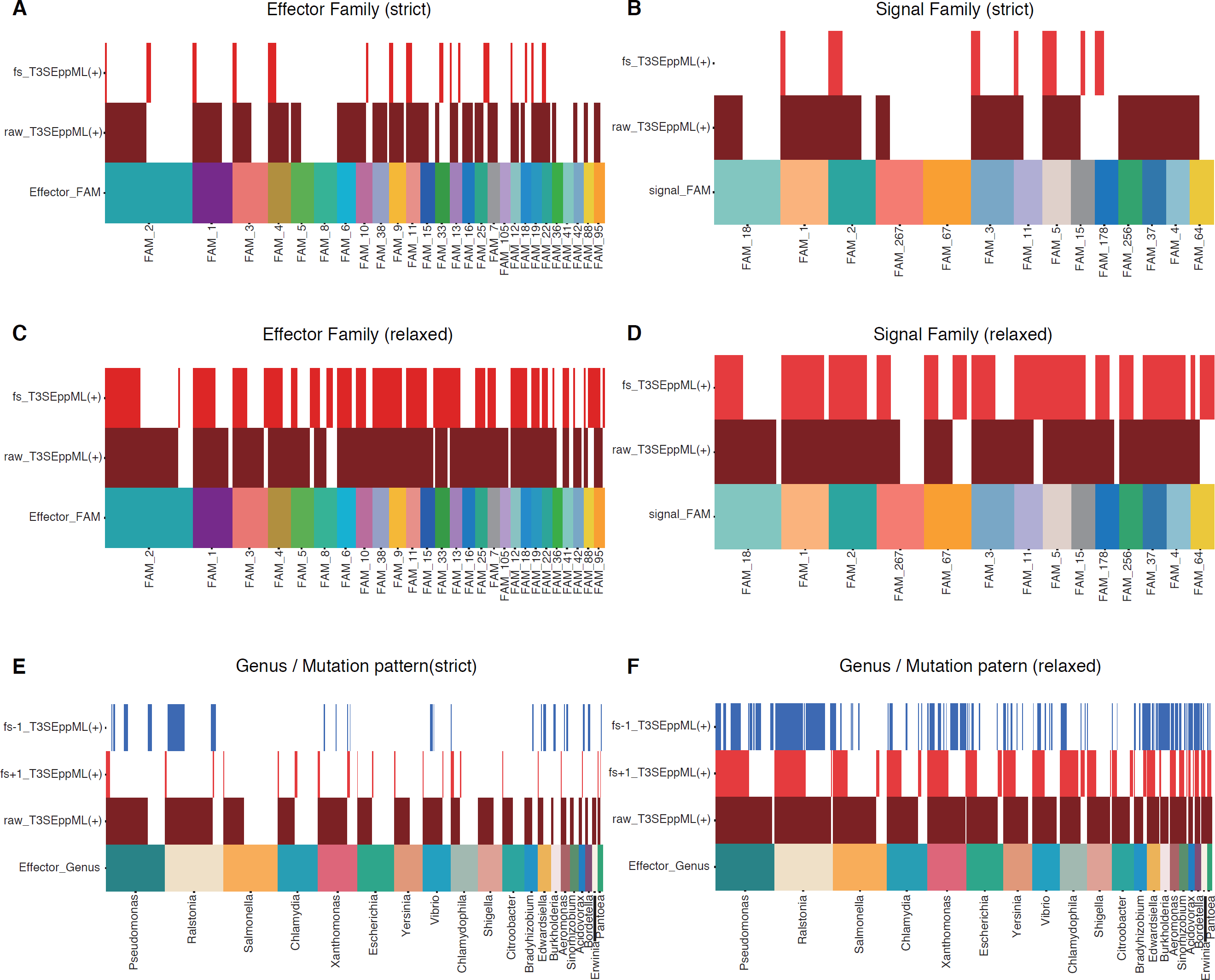
Distribution and preference of frameshift-tolerant effectors. The effector domain family (**A, C**) and signal family (**B, D**) distribution of T3SEppML prediction results were shown for the N-terminal 100-aa peptides of raw and frameshift-mutated type III effectors at the strict (=6) (**A, B**) and relaxed (>=3) (**C, D**) cutoffs. The genus and frameshift pattern distribution was also shown for the effectors at both the strict (**E**) and relaxed (**F**) cutoffs. The Effector families and Signal families referred to Hui et al., 2020. FAM, family; fs-1, “-1” pattern of frameshifts; fs+1, “+1” pattern of frameshifts.

When the cutoff was relaxed, 66.7% (248/372) and 67.5% (282/418) of the recalled effectors were tolerant to frameshifts, and they were distributed in 83.0% (83/100) of the covered effector families and 75.8% (191/252) of the signal families, respectively (Figure 2C-D; Supplemental Dataset S5). Among the large families, effector families such as Effector_FAM_11 (7/7), Effector_FAM_12 (5/5), Effector_FAM_13 (5/6), Effector_FAM_25 (5/6), Effector_FAM_33 (6/6) and Effector_FAM_38 (8/8), and signal families including SigFAM_1 (9/10), SigFAM_2 (8/10), SigFAM_5 (6/6) and SigFAM_37 (5/5), showed larger percentage of effectors tolerating frameshifts, despite the lack of statistical significance when compared to the overall frameshift-tolerating percentage (Supplemental Dataset S5). The effector family Effector_FAM_11 (XopG/NleD family distributed in both animal and plant pathogens), and signal families SigFAM_2 (being distributed in plant pathogens) and SigFAM_5 (in both animal and plant pathogens) were of particular interest, since they were relatively enriched at different cutoffs, for the effectors with tolerance of frameshifts (Supplemental Dataset S5).

Taken together, the results demonstrated that the type III effectors showing tolerance to frameshifts were widely distributed in various effector domain families or signal sequence families. There were, but only a few families, in which the effectors showed preference of frameshift tolerance.

### 4. Bacterial genus and mutation pattern preference of type III effectors tolerating frameshifts in mRNA encoding the N-terminal sequences

The type III effectors tolerant to frameshifts were also distributed in various bacteria, covering 60.9% (14/23) of the genera and 39.6% (19/48) of the species with raw effectors correctly recalled at the strict cutoff, and covering 92.3% (24/26) of the genera and 96.0% (48/50) of the species at the relaxed cutoff (Supplementary Dataset S5). At the relaxed cutoff, *Ralstonia* was the only genus for which the effectors tolerant to frameshifts were enriched marginally significantly compared to the distribution of effectors in other genera (63/69 vs. 309/474; EBT, *p* = 0.067; Figure 2E-F). The significance for *Ralstonia* became more striking when the strict cutoff was set (25/58 vs. 41/287; EBT, *p* = 0.019; Figure 2E-F). Besides *Ralstonia*, *Xanthomonas* (43/47), *Pseudomonas* (55/69), *Chlamydophila* (22/25), *Edwardsiella* (13/15), *Burkholderia* (*B. pseudomallei* specifically, 10/10), *Aeromonas* (10/11), *Acidovorax (A. citrulli* specifically, 6/6) also showed enrichment of the frameshift-tolerant effectors at the relaxed cutoff despite a lack of the significance, and *Pseudomonas* (13/51), *Chlamydophila* (4/11) and *Vibrio* (5/24) showed relatively larger proportion of frameshift-tolerant effectors than other genera at the strict cutoff (Figure 2E-F; Supplementary Dataset S5).

The frameshift patterns, “+1” and “-1”, also showed biases in some bacterial genera (Figure 2E-F; Supplementary Dataset S5). At the strict cutoff, most *Ralstonia* (23/25) effectors showed specific “-1” frameshift toleration (EBT, *p* = 1.94e-05). Although the proportion of “+1” frameshift-tolerant effectors increased when the cutoff was relaxed, most of which tolerated both “+1” and “-1” frameshifts in fact, the effectors of “-1”-pattern tolerance were still enriched significantly (60/63 vs. 38/63; EBT, *p* = 0.002). *Burkholderia* (10/10 vs. 1/10; EBT, *p* = 0.007) also showed preference of “-1” pattern, and *Chlamydophila* (22/22 vs. 7/22; EBT, *p* = 0.018), *Chlamydia* (18/20 vs. 7/20; EBT, *p* = 0.018) and *Shigella* (11/11 vs. 1/11; EBT, *p* = 0.004) showed preference of “+1” pattern, all at the relaxed cutoff (Figure 2E-F; Supplementary Dataset S5).

### 5. Common features shared by the N-terminal peptide sequences of raw and frameshifting effector proteins

We compared the basic sequence-derived statistical features of the raw and frameshift-mutated N-terminal signal sequences of type III secreted effectors, including sequential amino acid composition, bi-residue composition, one-hot encoding position-specific amino acid composition and physiochemical property binned amino acid composition.

The sequential composition of individual amino acids or bi-residues, or one-hot encoding position-specific amino acid composition profile could separate the raw and frameshifting N-terminal sequences of the effectors (Figure 3A-C). The frameshifting sequences predicted by T3SEppML as positive results (noted as “positive frameshifts”) were also clustered and closer to the raw sequences (Figure 3A-C). Physicochemical properties could cluster the “positive frameshifts” and the raw sequences closely, which were more clearly separated from the frameshifting sequences predicted by T3SEppML as negative signal sequences (noted as “negative frameshifts”) (Figure 3D). As controls, neither physicochemical properties nor amino acid composition profiles could well separate the N-terminal and C-terminal sequences of type III effectors (Figure 3E-F).

**Figure 3.**
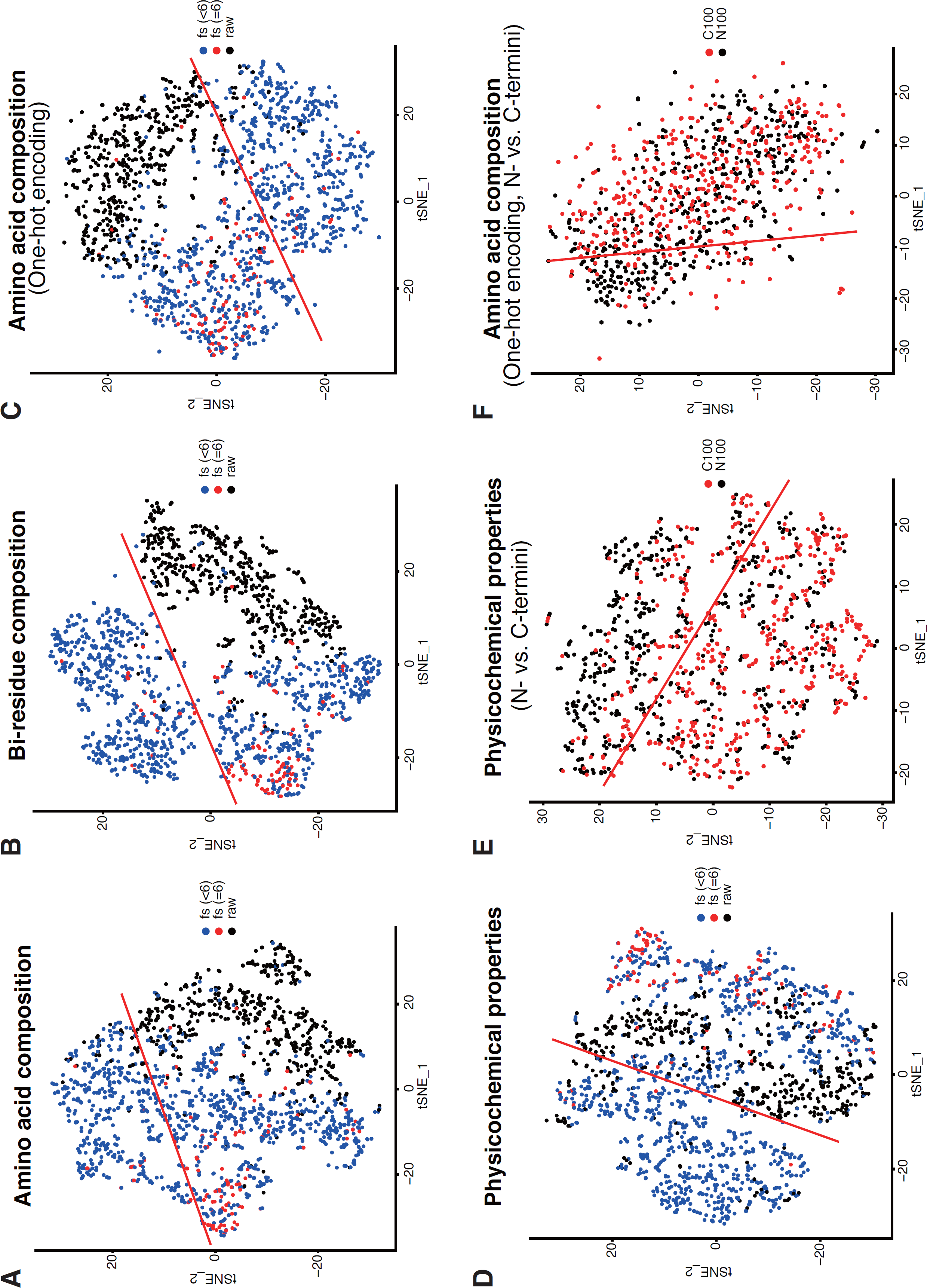
Clustering of the N-terminal 100-aa peptide sequences of raw and frame-shifted type III effectors. Sequential composition profile of individual amino acids (**A**), bi-residues (**B**) and physicochemical property binned amino acids (**D**), and one-hot encoding position-specific amino acid composition features (**C**) were used for clustering of the raw and frameshift-mutated sequences. Physicochemical features (**E**) and sequential amino acid composition features (**F**) were also used for clustering the control sequences, N-terminal and C-terminal 100-aa peptides of the raw type III effectors. In (**A**) - (**D**), the raw sequences, frameshift-mutated sequences showing tolerance (=6), and frameshift-mutated sequences without tolerance (<6) were shown in different colors. Red lines represented a decision line best separating the frameshift-mutated sequences without tolerance (<6) and the raw sequences or frameshift-mutated sequences showing tolerance (=6). In (**E**) and (**F**), the N-terminal and C-terminal peptide sequences were shown in different colors, and the decision line best separated the two types of sequences.

We also observed the distribution of raw and mutated sequences of type III effectors at the relaxed cutoff of T3SEppML. Apparently, a large number of the false positives according to the decision lines in Figure 3 became true positives, the “positive frameshifts” remained close to the raw effectors, and both the “positive frameshifts” and the raw effectors were clearly separated from the “negative frameshifts” (Supplementary Figure S1).

Therefore, for the effectors that tolerated frameshifts, the N-terminal sequences after frameshifting mutations retained common features with the raw sequences.

### 6. A self-correction mechanism of type III secretion signal sequences on frameshifts

Since *Ralstonia* showed larger percentage of effectors tolerant to frameshifts in N-terminal sequences, we aligned their protein and nucleotide sequences of tolerating effectors against sequence databases to observe if frameshift mutations happened in these genes naturally.

Generally, the genes could find mutated homologs in other strains or species, and the mutations happened more frequently in 5’-end segments nucleotide sequences (Supplementary Text S1). There were much fewer insertions or deletions (abbreviated as “indels”) than substitutions (Supplementary Text S1). An orphan indel, that is, a single indel in a relatively long region, >=60-nucleotide fragment for example, was often with nucleotides whose size could be divided by 3 with no remainder (Figure 4A). Therefore, the indels seldom caused sequence changes for long peptide fragments.

**Figure 4.**
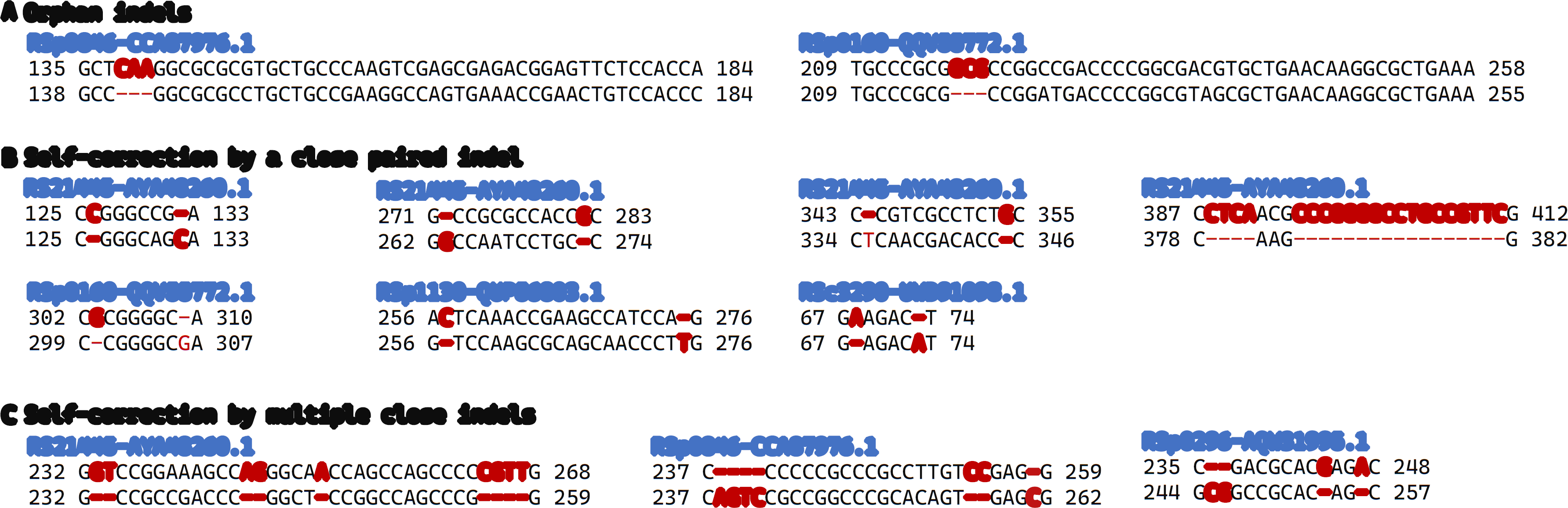
Natural indels present in the nucleotide sequences encoding the N-terminal peptides of *Ralstonia* type III effectors. (A) Examples of orphan indels. **(B)** Example of self-correction by a close paired indel. (C) Example of self-correction by multiple close indels.

However, indels of 3-individible nucleotides could still be identified in the nucleotide sequences encoding N-termini of type III effectors HrpF (RS_RS21445), skwp5 (Rsp0296), Rsp0846, RSc3290, Rsp0160 and Rsp1130 (Figure 4B). For these frameshifting mutations, paired indels were often identified nearby, normally with a <30-nucleotide span. The paired indels could complement the companion indels in that the summation of their introduced or missed nucleotides was the multiple of 3 and no longer caused frameshift (Figure 4B). Sometimes, multiple indels were required to block the continuing frameshift effect (Figure 4C).

Taken together, the results demonstrated that type III effectors could harness multiple strategies to avoid the possible influence of frameshift mutations on their effective secretion, such as self-correction by multi-locus complementary indels, or similar amino acid features before and after mutations.

### 7. Common mRNA features facilitating prediction of type III and IV effectors

The broad tolerance on frameshifting mutations in mRNA encoding the N-terminal of type III effectors (default settings for both T3SEppML and Bastion3), suggested that there could be common, albeit subtle, features buried in the mRNA level. Moreover, even though amino acid sequences were considered to bear the secretion signals for more type III or type IV effectors, the mRNA sequences encoding the putative signal peptide fragments could still show common features. To test this hypothesis, based on pre-trained transformers, we developed transferring learning models predicting type III and type IV effectors with mRNA sequences encoding the N-terminal and C-terminal 100 amino acids respectively, and observed the performance of the models.

For type III effectors, despite poorer performance compared to the models based on amino acid sequences, the DNABERT-based model, T3RNAtl, could still show a certain level of capability to classify type III effectors and non-effectors, with the Area Under the Receiver Operating Characteristic Curve (rocAUC) of 0.74 on an independent testing dataset (Figure 5A). We also tried other transformers pre-trained on nucleic acid sequences. The models based on Routing Transformer and Transformer also showed some capability in predicting type III effectors with rocAUC of 0.70 and 0.66 respectively (Figure 5A). Therefore, the mRNA sequences encoding the N-termini of type III effectors also contained some common features. However, the classification performance was generally not good, and when an optimal cutoff was selected, the specificity and sensitivity achieved 0.79 and 0.58 respectively (Figure 5B).

**Figure 5.**
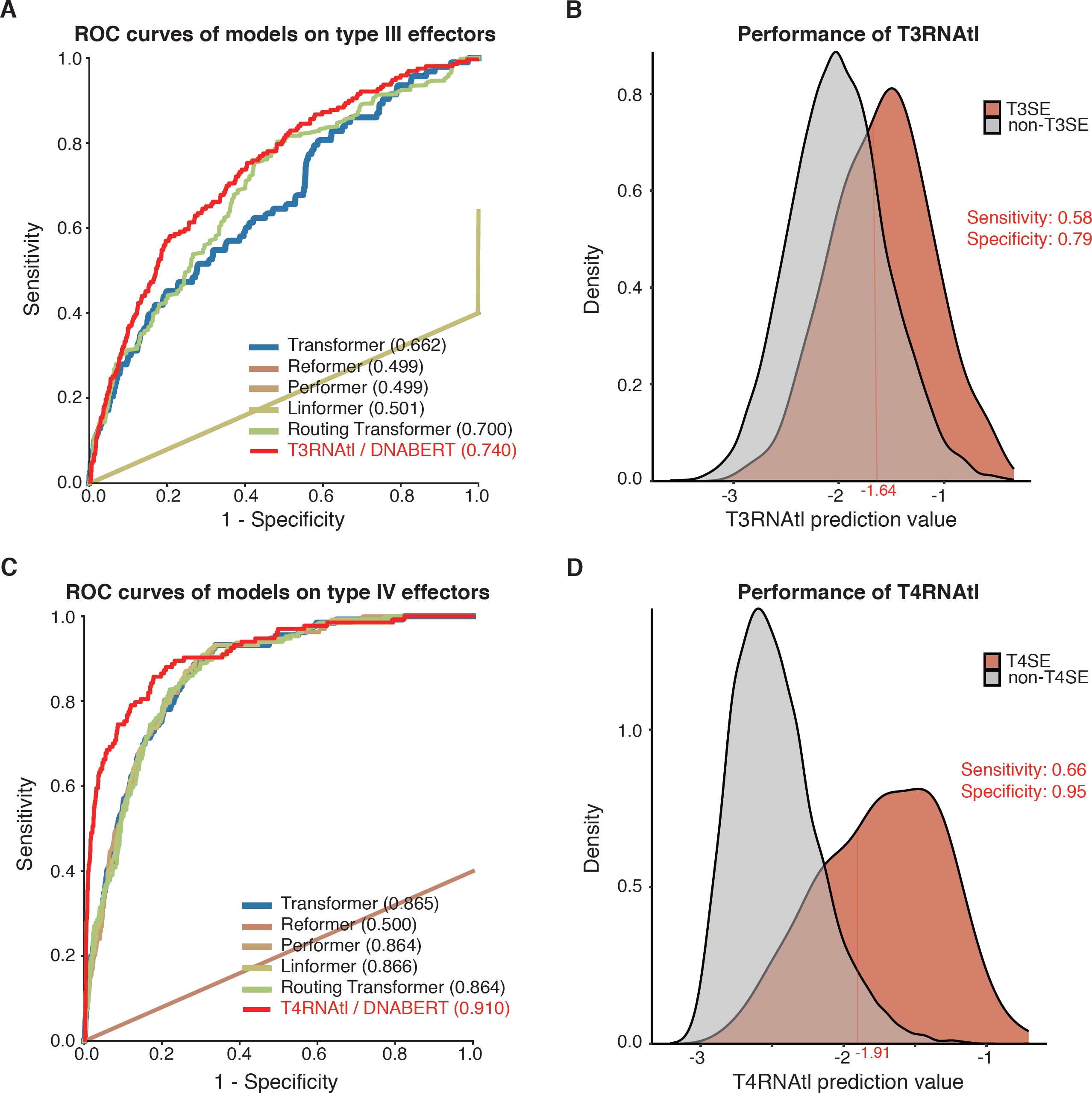
Performance of the natural language processing models predicting type III and IV effectors using mRNA sequences. (**A**) ROC curves of transformer-based models predicting the testing dataset of type III effectors and negative sequences. (**B**) Distribution density curves of the T3RNAtl predicting values on the positive and negative sequences in the testing dataset of type III effectors and negative sequences. **(C)** ROC curves of transformer-based models predicting the testing dataset of type IV effectors and negative sequences. (**D**) Distribution density curves of the T4RNAtl predicting values on the positive and negative sequences in the testing dataset of type IV effectors and negative sequences. In (**B**) and (**D**), An optimized decision cutoff was selected for T3RNAtl and T4RNAtl and indicated respectively, according to which the specificity and sensitivity was calculated and shown.

The transformer-based models were also trained for the mRNA sequences encoding the C-terminal peptides of type IV effectors and negative control proteins. Not like the type III effectors, the models showed good performance in predicting type IV effectors, with T4RNAtl based on DNABERT performing best, followed by Linformer, Transformer and Performer (Figure 5C). T4RNAtl reached a rocAUC of 0.91 (Figure 5C). The specificity and sensitivity could reach 0.95 and 0.66 respectively, when the cutoff was optimized (Figure 5D). Therefore, type IV effectors showed more typical, common features in the mRNA sequences encoding their C-terminal peptides.

## Discussion

Previous studies demonstrated that the mRNA encoding N-terminal peptides of type III effectors showed tolerance to frameshifts (Anderson and Schneewind 1997; Mudgett et al., 2000; Rüssmann et al., 2002; Arnold et al., 2009; Wang et al., 2011). However, only a few effectors were shown with the capability. Here, with a machine learning approach (T3SEppML), we found that many more type III effectors could tolerate frameshifts in mRNA encoding the N-termini. It was not likely due to high false positive rate of T3SEppML, since similar observation was obtained with other state-of-the-art prediction tools, and when the specificity was increased, the percentage of frameshift-tolerant type III effectors remained high, significantly larger than that of C-termini of type III effectors or the signal fragments of type IV, type I and Sec/Tat secreted substrates. It was noted that, when the strict cutoff was selected, there were a lot of type III effectors that were predicted to be non-effectors for the raw sequences but predicted to be effectors after frameshifting mutations (Figure 1D). It strengthened the generality and capacity of type III effectors in tolerance against frameshifts, though this subset of effectors were not included for further analysis. We also noted that the C-termini of type III effectors showed larger proportion than the signal peptides of type IV, type I or Sec/Tat secreted substrates in tolerance of mRNA frameshifts (Figure 1E-H). It was not surprising, since previous studies demonstrated that secretion signals could also be presented in the C-termini of type III effectors (Tomalka et al., 2012; Login and Wolf-Watz 2015).

Frameshift tolerance of secretion signals was observed in type III effectors belonging to various effector families, signal peptide families, and bacterial species or genera. Therefore, it is a general phenomenon in type III effectors. However, the frameshift-tolerant effectors also showed preference in some effector and signal families, as could be related with the specific sequence properties of these effectors (Supplementary Dataset S5). Some bacteria, especially *Ralstonia*, showed significantly larger proportion of effectors tolerant to frameshift mutations (Supplementary Dataset S5). Moreover, in many of these or other genera or species, the effectors showed striking preference to a certain pattern of frameshifts (“+1” or “-1”) (Supplementary Dataset S5). It remains to be clarified about the mechanisms for such preferences, which could be related with specific evolution of bacteria and/or their type III secretion systems or effectors.

It was shown that, the sequences encoding N-terminal peptides of type III effectors, experienced strong positive selection (Sato et al., 2011). We also found that the mutations happened in this region more frequently than in the region encoding effector domains (Supplementary Text S1). Not limited to substitutions, indels happened too. However, by self-correction mechanisms, the indels seldom influenced a long span of amino acids naturally. Sometimes, a short stretch of amino acids, for example, the residues 78-89 in *Ralstonia* RS21445 (Figure 4C), could still be involved. Even if the frameshifts influenced longer peptides, the changed peptides could retain the overall similar composition profiles and physicochemical properties of amino acids in the raw sequences (Figure 3; Supplementary Figure S1). Therefore, bacteria manipulate multiple tactics to avoid likely negative influence in recognition of type III secretion caused by frameshift mutations. The peptide sequences could potentially still play critical roles in guiding the specific secretion of most type III effectors. The mRNA sequences encoding N-terminal peptides also exhibited common features among type III effectors, which were different from those of non-effectors (Figure 5A-B). However, the features are very subtle, possibly encoding the atypical common features in amino acid sequences. Compared to genes encoding type IV effectors and other secreted proteins, terminal reassortment was found to happen frequently and specifically in type III effector genes (Stavrinides et al., 2006). The terminal recombination events could possibly introduce indels and frameshift mutations. Consequently, bacteria have evolved strategies to remedy the deficiencies in type III secretion and the function of effectors. The strategies might include positively selected mutations, paired self-corrected indels, and retaining the physicochemical properties or other key features of raw signal peptide sequences.

Compared to type III effectors, the signal sequences of type IV effectors appeared more conserved in nucleotide level (Figure 5C-D). Frameshifts seldom happened in the signal regions of type IV effectors although they derived peptide sequences most of which were predicted no long to contain type IV secretion signals (Figure 1F). Therefore, it is also likely that conserved properties in mRNA sequences encode common features of amino acids in peptide sequences that are important for specific recognition of type IV secretion. Domain reshuffling has also been widely disclosed in type IV effectors (Burstein et al., 2016; Gomez-Valero et al., 2019), but it could exert little influence on C-terminal peptides of the proteins by introduction of indels in the nucleotide sequences via local recombination. Moreover, the sequences encoding C-termini of proteins should experience much stronger purifying selection pressure for frameshift mutations since there would be much smaller space for remedy. Therefore, the type IV effectors do not have to evolve strategies to tolerate the frameshifts in the mRNA sequences encoding the C-terminal signal peptides.

In this study, we found an unexpected number of type III effectors tolerant to the frameshift mutations in mRNA encoding the N-terminal peptide sequences. The frameshift tolerance of secretion signals was not found in type IV, type I and Sec/Tat secreted proteins. The type III effectors with frameshift tolerance of secretion signals were distributed in various effector families, signal peptide families and bacterial genera or species, but also showed preference. The natural frameshifts were often remedied by paired indels in the sequences encoding the N-terminal peptides of type III effectors. Frameshift-derived peptide sequences also retained the common properties present in the signal peptides of raw type III effectors. Finally, common features could be found in the mRNA sequences encoding N-terminal 100-aa peptides of type III effectors and those encoding C-terminal 100-aa peptides of type IV effectors, with which natural language processing models could be trained, to predict type III and type IV effectors, respectively.

## Materials and Methods

### 1. Datasets

The protein and nucleotide sequences for 618 experimentally verified type III effectors, 407 type IV effectors and 93 type I secreted proteins were downloaded from TxSEdb (http://61.160.194.165:3080/TxSEdb). The genome sequences and annotation files for *Acinetobacter baumannii* K09-14 (Genome Accession: CP043953.1), *Advenella kashmirensis* WT001 (Genome Accession: NC_017964.1), *Agrobacterium tumefaciens* 12D1 (Genome Accession: CP033031.1 and CP033032.1), *Klebsiella pneumonia subsp. pneumoniae* HS11286 (Genome Accession: CP003200.1), *Stenotrophomonas* sp. CD2 (Genome Accession: CP102248.1), *E. coli* MG1655 (Genome Accession: NC_000913.3), *S.* typhimurium 14028S (Genome Accession: NC_016856.1), *Y. entericolitica* 8081 (Genome Accession: NC_008800.1), and *R. solanacearum* GMI1000 (Genome Accession: NC_003296.1) were downloaded from NCBI Genome database (http://www.ncbi.nlm.nih.gov/genome).

### 2. *In silico* modeling of frameshift mutations

Both “+1” and “-1” frameshift mutations were introduced to genes encoding the type III effectors, type IV effectors, type I secreted proteins, proteins of *E. coli* MG1655, *S. typhimurium* 14028S, *Y. entericolitica* 8081 and *R. solanacearum* GMI1000. For “+1” frameshifts, an adenosine was inserted immediately after the starting codon while the nucleotide preceding the stop codon was removed. For “-1” frameshifts, the first nucleotide immediately after the starting codon was removed while an adenosine was added before the stop codon. The mutated nucleotide sequences were translated into proteins sequences with classical codons, except for replacing each premature stop codon with an alanine.

### 3. Tolerance of frameshift mutations in mRNA sequences encoding the signal peptides

The N-terminal 100-aa peptides of raw type III effectors and the framshift-derived protein sequences were predicted with the T3SEppML module integrated in T3SEpp, with the default relaxed cutoff (>=3/6) or a strict cutoff (=6) (Hui et al., 2020). The peptide sequences of C-terminal 100-aa peptides derived from the raw type III effectors were also predicted with T3SEppML at the strict cutoff. Another tool, Bastion3, which by default predict type III effectors with the full-length protein sequences, was also used to predict the raw and frameshift-mutated N-terminal 100-aa sequences of type III effectors (Wang et al., 2019). Default settings were used for Bastion3.

The C-terminal 100-aa peptides were retrieved from the raw and frameshift-mutated type IV effectors, and predicted by T4SEpre with a cutoff of 3/3, that is, positive prediction by all the three modules, T4SEpre_bpbaac, T4SEpre_pseaac and T4SEpre_joint (Wang et al., 2014). T1SEstacker was used to predict the C-terminal 60-aa fragments of the raw and frameshift-mutated type I secreted proteins, with a default cutoff (>=3/6) (Chen et al., 2022). For the Sec/Tat signal peptides, we used SignalP 6.0 and the default settings to screen all possible Sec/Tat substrates from the raw and frameshift-mutated proteome of *E. coli* MG1655, *S. typhimurium* 14028S, *Y. entericolitica* 8081 and *R. solanacearum* GMI1000 (Teufel et al., 2022).

### 4. Distribution analysis of the type III effectors tolerant to frameshifts

The signal families and effector domain families of type III effectors referred to the previous study (Hui et al., 2020). The annotation information for the bacterial genus and species of each type III effector was downloaded from TxSEdb (http://61.160.194.165:3080/TxSEdb). The composition rates were compared with EBT, a statistic test balancing power and precision (Hui et al., 2017). The significance level was preset as *p* < 0.05.

### 5. Amino acid composition and physicochemical properties of the N-terminal peptides of raw and frameshift-mutated type III effectors

For each of the N-terminal 100-aa peptides from the raw and frameshift-mutated type III effectors, the sequential composition of individual amino acids (20 features) and continuous bi-residues (400 features) were calculated, respectively. Each 100-aa sequence was also represented using one-hot encoding scheme for the amino acid in each position (20-bit “0/1” features for the amino acids in each position; in total 2000 features). The amino acids in each sequence were also binned according to nine physicochemical properties (tiny, small, aliphatic, aromatic, polar, non-polar, charged, basic and acidic), and then sequential composition of amino acids for each property was calculated (9 features) (Hobbs et al., 2016). Each sequence was represented as a vector of the sequential amino acid composition features, bi-residue composition features, position-specific one-hot encoding features and sequential composition of physicochemical property binned amino acids, respectively. All the sequences represented by a *n * m* matrix for each type of features, where *n* and *m* meant the number of sequences and the dimension of features respectively, were clustered and visualized by a t-Distributed Stochastic Neighbor Embedding (t-SNE) method (van der Maaten and Hinton 2008).

### 6. Detection of frameshift mutations happening naturally in genes encoding type III effectors

The protein sequences of 23 *R. solanacearum* GMI1000 effectors showing tolerance to frameshift mutations at the strict cutoff (Supplementary Dataset S5) were aligned against other *Ralstonia* proteins curated in NCBI non-redundant (NR) protein database using Protein BLAST (https://blast.ncbi.nlm.nih.gov/Blast.cgi). The hits without 100% identity were checked manually and the representatives with amino acid indels were retrieved. The nucleotide sequences encoding the paired proteins were also retrieved and pairwise alignment was performed using the Nucleotide BLAST program.

### 7. Natural language processing models predicting type III and type IV effectors with mRNA sequences

The nucleotide sequences encoding the N-terminal 100-aa peptides of verified type III and type IV effectors were used as the raw positive dataset. We selected the whole proteomes of representative bacterial strains without a type III secretion system (*A. baumannii* K09-14, *A. kashmirensis* WT001, *A. tumefaciens* 12D1, *K. pneumoniae subsp. pneumoniae* HS11286, *Stenotrophomonas sp*. CD2 and *E. coli* MG1655) or a protein-translocating type IV secretion system (*A. baumannii* K09-14, *A. kashmirensis* WT001, *K. pneumoniae subsp. pneumoniae* HS11286, *Stenotrophomonas sp.* CD2 and *E. coli* MG1655) as the origin of raw negative dataset for type III and type IV effectors, respectively, and the 300-nucleotide fragments were extracted from the genes encoding the N-terminal 100-aa peptides. CD-HIT was used to remove the homolog-redundant sequences from the positive dataset, the negative dataset, and between the positive and negative datasets, with a cutoff similarity of 0.6 (Li and Godzik 2006). After the homology filtering, 542 positive type III sequences (T3SE) and 383 positive type IV sequences (T4SE) remained, and 24085 negative type III sequences (non-T3SE) and 19567 negative type IV sequences (non-T4SE) were retained. All the positive and negative datasets were separated randomly into two parts as a 2:1 ratio, with 2/3 of the sequences as the training datasets and the remaining 1/3 of the sequences as the testing datasets. In the training datasets, the proportion was ∼2.25% for type III effectors and ∼1.96% for type IV effectors, approaching the real distribution of type III and type IV effectors in most bacterial strains bearing a type III or type IV secretion system.

The DNABERT pre-trained transformer was downloaded, and was used to extract context information from input sequences (Ji et al., 2021). Transfer learning models were trained using the DNABERT-generated features to predict type III effectors (T3RNAtl) and type IV effectors (T4RNAtl) respectively. The transfer learning models were Deep Neural Network (DNN) based, with two fully connected layers of 1024 and 512 nodes respectively. The entire models were fine-tuned using the hyperparameters including: optimizer, SGD (Stochastic Gradient Descent), optimizer momentum as 0.9, and optimizer learning rate as 0.1. Each model was trained for a total of 4 epochs. Transfer learning models based on other DNA/RNA pre-trained transformers were also trained and evaluated using the online service of DeepBIO with the default settings (Wang et al., 2023). The other pre-trained transformers used for prediction of type III and type IV effectors included Transformer, Reformer, Performer, Linformer and Routing Transformer.

The performance of models was evaluated using the testing datasets as mentioned above. ROC curves were not influenced by the selection of decision cutoff and therefore used as the fair measure of the model performance. The AUC was calculated for each model. An optimized cutoff was selected for T3RNAtl and T4RNAtl respectively. Accordingly, sensitivity and specificity were calculated as “true positive predictions / all positive sequences” and “true negative predictions / all negative sequences” respectively.

### 8. Codes accessibility

The codes for T3RNAtl and T4RNAtl were accessible freely through the links: http://61.160.194.165:3080/TxRNA, or https://github.com/Bricker-Jimmy/TxRNAdl.

### 9. Statistics

The statistic tests involved in the study were described in context. For all the tests, the threshold for type I error was preset as α < 0.05.

## Contribution of Authors

Y.W. conceived the project. J.Y. and M.L. supervised the project and conducted most of the analysis. M.Y., X.W., L.L., X.C., R.C. collected, annotated the effector sequences and conducted the modelling analysis. J.Y., Z.Z., R.C. and Y.H. trained the machine learning models. J.Y., M.L. and Y.W. wrote the first draft of manuscript. All the authors revised the manuscript and approved the final version.

## Acknowledgements

The work was supported by a Natural Science Fund of Shenzhen (JCYJ20190808165205582) to Y.W., and by a National Undergraduate Training Program for Innovation and Entrepreneurship (no. 202210590048) to X.C.

## Supplementary Materials

**Supplementary Text S1. Examples of indels naturally happening in the nucleotide sequences encoding the N-terminal peptides of Ralstonia type III effectors.**

**Supplementary Figure S1. Clustering of the N-terminal 100-aa peptide sequences of raw and frame-shifted type III effectors.** Same with Figure 3 except that the cutoff for toleration of frameshift mutation was set as >=3.

**Supplementary Dataset S1. Prediction results of T3SEppML or Bastion3 on N-terminal or C-terminal 100-aa peptides from the raw or frameshift-mutated type III effectors.**

**Supplementary Dataset S2. Prediction results of T4SEpre on C-terminal 100-aa peptides from the raw or frameshift-mutated type IV effectors.**

**Supplementary Dataset S3. Prediction results of T1SEstacker on C-terminal 60-aa peptides from the raw or frameshift-mutated type I secreted proteins.**

**Supplementary Dataset S4. Prediction results of SignalP 6.0 on the raw or frameshift-mutated potential Sec/Tat substrate proteins in representative bacterial strains.**

**Supplementary Dataset S5. Distribution and preference of frameshift tolerant type III effectors.**

## References

Anderson DM, Fouts DE, Collmer A, Schneewind O. (1999a). Reciprocal secretion of proteins by the bacterial type III machines of plant and animal pathogens suggests universal recognition of mRNA targeting signals. Proc Natl Acad Sci USA. 96:12839–12843.

Anderson DM, Schneewind O. (1997). A mRNA signal for the type III secretion of Yop proteins by Yersinia enterocolitica. Science. 278:1140–1143.

Anderson DM, Schneewind O. (1999b). Yersinia enterocolitica type III secretion: an mRNA signal that couples translation and secretion of YopQ. Mol Microbiol 31: 1139–1148.

Arnold R, Brandmaier S, Kleine F, Tischler P, Heinz E, Behrens S, Niinikoski A, Mewes HW, Horn M, Rattei T. (2009). Sequence-based prediction of type III secreted proteins. PLoS Pathog. 5(4):e1000376.

Arnold R, Jehl A, Rattei T. (2010). Targeting effectors: the molecular recognition of Type III secreted proteins. Microbes Infect. 12:346–358.

Bartonek L, Braun D, Zagrovic B. (2020). Frameshifting preserves key physicochemical properties of proteins. Proc Natl Acad Sci U S A. 117(11):5907–5912.

Blaylock B, Sorg JA, Schneewind O. (2008).Yersinia enterocolitica type III secretion of YopR requires a structure in its mRNA. Mol Microbiol. 70: 1210–1222.

Burstein D, Amaro F, Zusman T, Lifshitz Z, Cohen O, Gilbert JA, Pupko T, Shuman HA, Segal G. (2016). Genomic analysis of 38 Legionella species identifies large and diverse effector repertoires. Nat Genet. 48(2): 167–175.

Chen Z, Zhao Z, Hui X, Zhang J, Hu Y, Chen R, Cai X, Hu Y, Wang Y. (2022). T1SEstacker: A Tri-Layer Stacking Model Effectively Predicts Bacterial Type 1 Secreted Proteins Based on C-Terminal Non-repeats-in-Toxin-Motif Sequence Features. Front Microbiol. 12:813094.

Fei J, Sharma CM. (2018). RNA Localization in Bacteria. Microbiol Spectrum. 6(5):RWR-0024-2018.

Galán JE, Waksman G. (2018). Protein-Injection Machines in Bacteria. Cell. 172(6):1306–1318.

Gomez-Valero L, Rusniok C, Carson D, Mondino S, Pérez-Cobas AE, Rolando M, Pasricha S, Reuter S, Demirtas J, Crumbach J, Descorps-Declere S, Hartland EL, Jarraud S, Dougan G, Schroeder GN, Frankel G, Buchrieser C. (2019). More than 18,000 effectors in the Legionella genus genome provide multiple, independent combinations for replication in human cells. Proc Natl Acad Sci U S A. 116(6):2265–2273.

Hu Y, Huang H, Cheng X, Shu X, White AP, Stavrinides J, Köster W, Zhu G, Zhao Z, Wang Y. (2017). A global survey of bacterial type III secretion systems and their effectors. Environ Microbiol. 19(10):3879–3895.

Hui X, Chen Z, Lin M, Zhang J, Hu Y, Zeng Y, Cheng X, Ou-Yang L, Sun MA, White AP, Wang Y. (2020). T3SEpp: an Integrated Prediction Pipeline for Bacterial Type III Secreted Effectors. mSystems. 5(4):e00288–20.

Hui X, Hu Y, Sun MA, Shu X, Han R, Ge Q, Wang Y. (2017). EBT: a statistic test identifying moderate size of significant features with balanced power and precision for genome-wide rate comparisons. Bioinformatics. 33(17):2631–2641.

Ji Y, Zhou Z, Liu H, Davuluri RV. (2021). DNABERT: pre-trained Bidirectional Encoder Representations from Transformers model for DNA-language in genome. Bioinformatics. 37(15):2112–2120.

Jing R, Wen T, Liao C, Xue L, Liu F, Yu L, Luo J. (2021). DeepT3 2.0: improving type III secreted effector predictions by an integrative deep learning framework. NAR Genom Bioinform. 3(4):lqab086.

Kannaiah S, Amster-Choder O. (2014). Protein targeting via mRNA in bacteria. Biochim Biophys Acta. 1843(8):1457–65.

Karavolos MH, Roe AJ, Wilson M, Henderson J, Lee JJ, Gally DL, Khan CM. (2005). Type III secretion of the Salmonella effector protein SopE is mediated via an N-terminal amino acid signal and not an mRNA sequence. J Bacteriol. 187(5):1559–67.

Li W, Godzik A. (2006). Cd-hit: a fast program for clustering and comparing large sets of protein or nucleotide sequences. Bioinformatics. 22(13):1658–9.

Lloyd SA, Norman M, Rosqvist R, Wolf-Watz H. (2001). Yersinia YopE is targeted for type III secretion by N-terminal, not mRNA, signals. Mol Microbiol. 39:520–531.

Lloyd SA, Sjöström M, Andersson S, Wolf-Watz H. (2002). Molecular characterization of type III secretion signals via analysis of synthetic N-terminal amino acid sequences. Mol Microbiol. 43:51–59.

Login FH, Wolf-Watz H. (2015). YscU/FlhB of Yersinia pseudotuberculosis harbors a C-terminal type III secretion signal. J Biol Chem. 290(43):26282–26291.

Majander K, Anton L, Antikainen J, Lång H, Brummer M, Korhonen TK, Westerlund-Wikström B. 2005. Extracellular secretion of polypep-tides using a modified Escherichia coli flagellar secretion apparatus. Nat Biotechnol. 23:475– 481.

Mudgett MB, Chesnokova O, Dahlbeck D, Clark ET, Rossier O, et al. (2000). Molecular signals required for type III secretion and translocation of the Xanthomonas campestris AvrBs2 protein to pepper plants. Proc Natl Acad Sci USA. 97: 13324–13329.

Nagai H, Cambronne ED, Kagan JC, Amor JC, Kahn RA, Roy CR. (2005). A C-terminal translocation signal required for Dot/Icm-dependent delivery of the Legionella RalF protein to host cells. Proc Natl Acad Sci USA. 102:826–831.

Nevo-Dinur K, Nussbaum-Shochat A, Ben-Yehuda S, Amster-Choder O. (2011). Translation-independent localization of mRNA in E. coli. Science. 331(6020):1081–1084.

Niemann GS, Brown RN, Mushamiri IT, Nguyen NT, Taiwo R, Stufkens A, Smith RD, Adkins JN, McDermott JE, Heffron F. (2013). RNA type III secretion signals that require Hfq, J Bacteriol. 195:2119–2125.

Noroy C, Lefranc, ois T, Meyer DF. (2019). Searching algorithm for Type IV effector proteins (S4TE) 2.0: Improved tools for Type IV effector prediction, analysis and comparison in proteobacteria. PLoS Comput Biol. 15(3): e1006847.

Ramamurthi KS, Schneewind O. (2003). Yersinia yopQ mRNA encodes a bipartite type III secretion signal in the first 15 codons. Mol Microbiol. 50(4):1189–98.

Rüssmann H, Kubori T, Sauer J, Galán JE. (2002). Molecular and functional analysis of the type III secretion signal of the Salmonella enterica InvJ protein. Mol Microbiol. 46:769–779.

Sato Y, Takaya A, Yamamoto T. (2011). Meta-analytic approach to the accurate prediction of secreted virulence effectors in gram-negative bacteria. BMC Bioinformatics. 12:442

Teufel F, Almagro Armenteros JJ, Johansen AR, Gíslason MH, Pihl SI, Tsirigos KD, Winther O, Brunak S, von Heijne G, Nielsen H. (2022). SignalP 6.0 predicts all five types of signal peptides using protein language models. Nat Biotechnol. 40(7):1023–1025.

Tomalka AG, Stopford CM, Lee PC, Rietsch A. (2012). A translocator-specific export signal establishes the translocator-effector secretion hierarchy that is important for type III secretion system function. Mol Microbiol. 86(6):1464–1481.

Van der Maaten L, Hinton G. (2008). Visulizing data using t-SNE. J Mach Learn Res. 9:2579–2605.

Wagner N, Avram O, Gold-Binshtok D, Zerah B, Teper D, Pupko T. (2022). Effectidor: an automated machine-learning-based web server for the prediction of type-III secretion system effectors. Bioinformatics. 38(8):2341–2343.

Wang J, Li J, Yang B, Xie R, Marquez-Lago TT, Leier A, Hayashida M, Akutsu T, Zhang Y, Chou KC, Selkrig J, Zhou T, Song J, Lithgow T. (2019). Bastion3: a two-layer ensemble predictor of type III secreted effectors. Bioinformatics. 35(12):2017–2028.

Wang R, Jiang Y, Jin J, Yin C, Yu H, Wang F, Feng J, Su R, Nakai K, Zou Q, Wei L. (2023). DeepBIO: an automated and interpretable deep-learning platform for high-throughput biological sequence prediction, functional annotation and visualization analysis. Nucleic Acids Res. gkad055.

Wang X, Dong Q, Chen G, Zhang J, Liu Y, Cai Y. (2022). Frameshift and wild-type proteins are often highly similar because the genetic code and genomes were optimized for frameshift tolerance. BMC Genomics. 23(1):416.

Wang Y, Wei X, Bao H, Liu SL. (2014). Prediction of bacterial type IV secreted effectors by C-terminal features. BMC Genomics. 15:50.

Wang Y, Zhang Q, Sun MA, Guo D. (2011). High-accuracy prediction of bacterial type III secreted effectors based on position-specific amino acid composition profiles. Bioinformatics. 27(6):777–784.

